## Supplemental figures for "Proteome efficiency of metabolic pathways in *Escherichia coli* increases along the nutrient flow"

This PDF includes Figure S1-S7 and supplementary references 1-13.

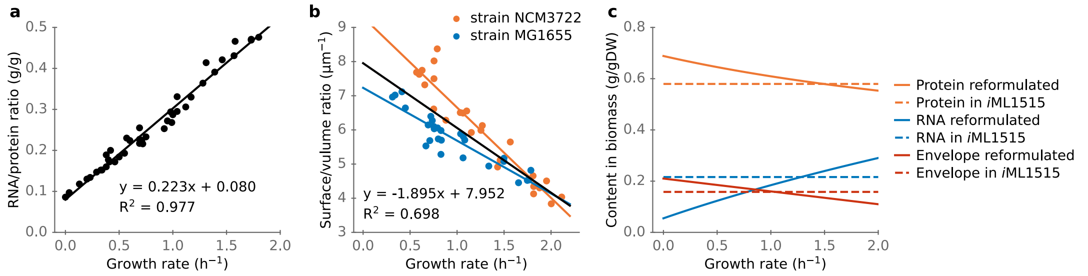


**Fig. S1. Growth rate-dependent biomass composition.** (**a**) Growth rate-dependent RNA/protein mass ratio; data from Ref.(1, 2). (**b**) Growth rate-dependent surface/volume ratio; data for wildtype strain in non-stressed growth conditions from Ref.(3). Surface/volume ratios are slightly different between the NCM3722 and MG1655 strains; to estimate the required production of cell envelope components, we used a regression on data from both strains (black line). **(c)** Growth rate-dependent biomass composition, reformulated considering the growth rate-dependent RNA/protein ratio in panel (a) and the surface/volume ratio in panel (b). The category "Envelope" includes murein, lipopolysaccharides, and lipids.

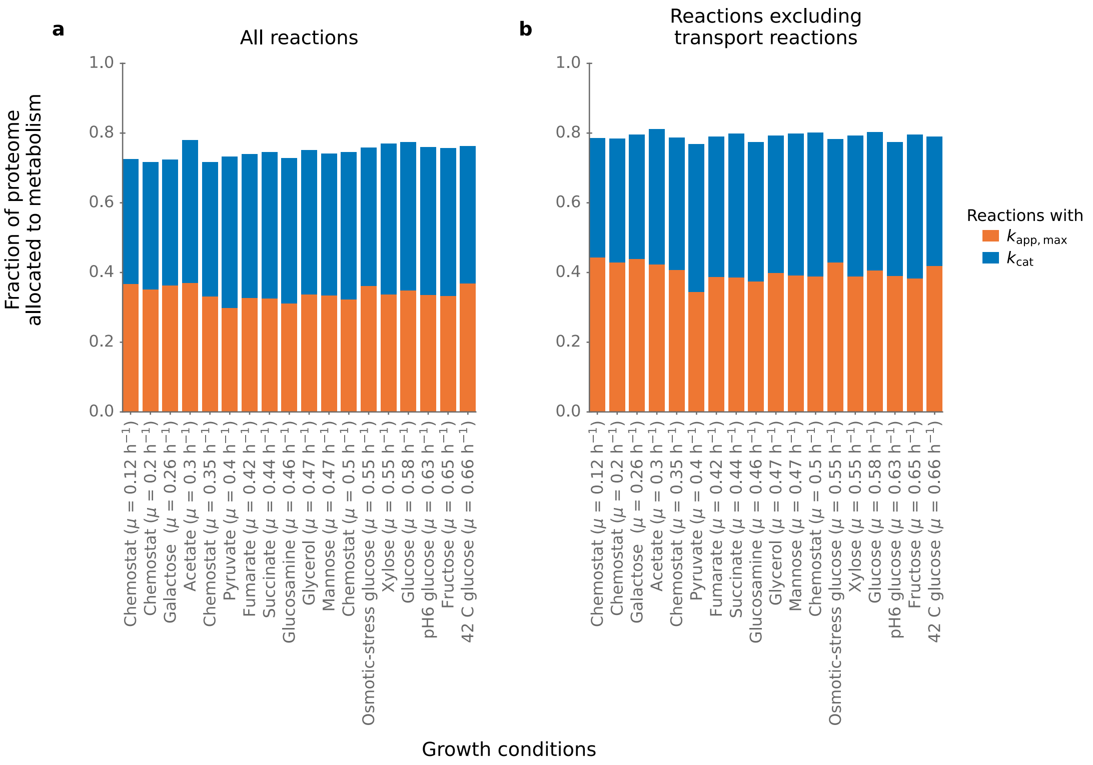


**Fig. S2. Reactions with available enzyme turnover numbers cover more than 70% of the proteome fraction allocated to metabolism.** Colors show the proteome fractions of enzymes with *in vivo* enzyme turnover number ($k_{app,max}$, in orange) or *in vitro* turnover number ($k_{kcat}$, in blue). (**a**) All reactions in iML1515. (**b**) All non-transport reactions in iML1515.

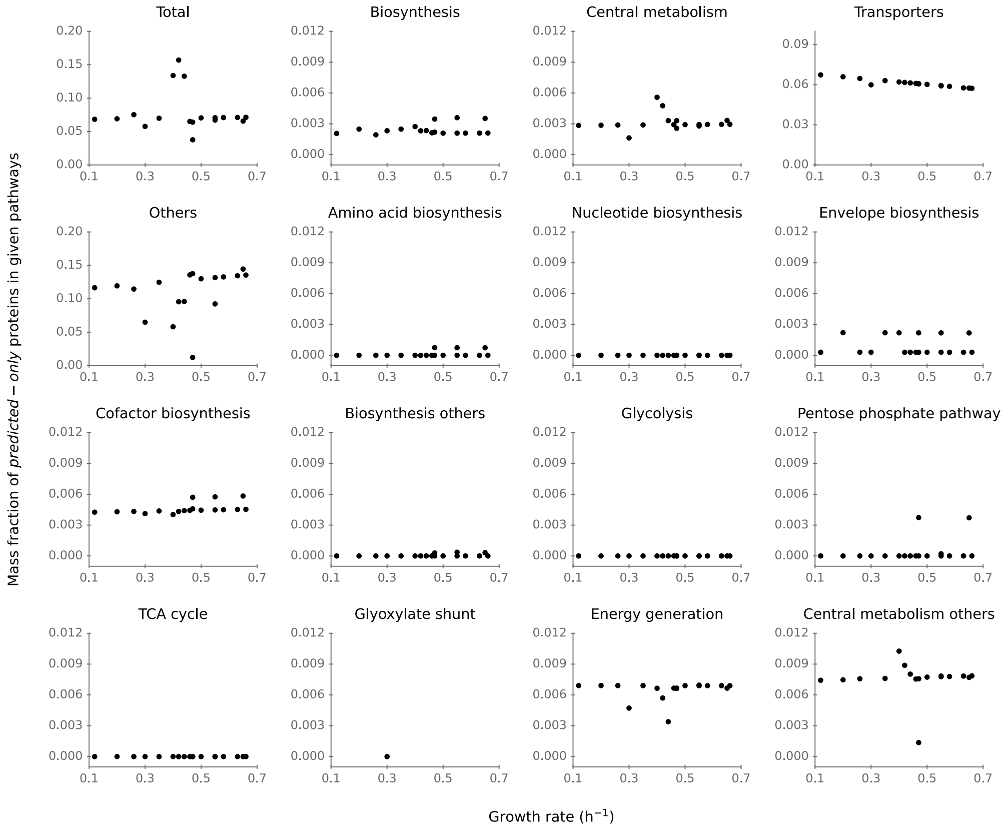


**Fig. S3. The mass fraction of predicted-only proteins across pathways is generally low.** Except for transporters and for proteins that cannot be assigned to particular pathways (Others), the *predicted-only* proteins account for less than 1% of the predicted proteins in each pathway.

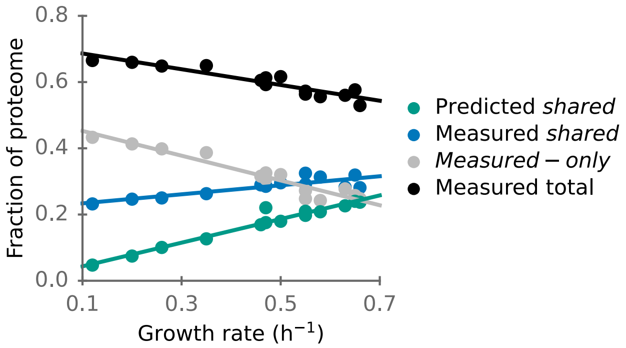


**Fig. S4. Proteome efficiency of the whole metabolic network.** With increasing growth rate, predicted and experimentally observed *shared* metabolic proteome fractions increase, while the *measured-only* metabolic proteome fraction decreases.

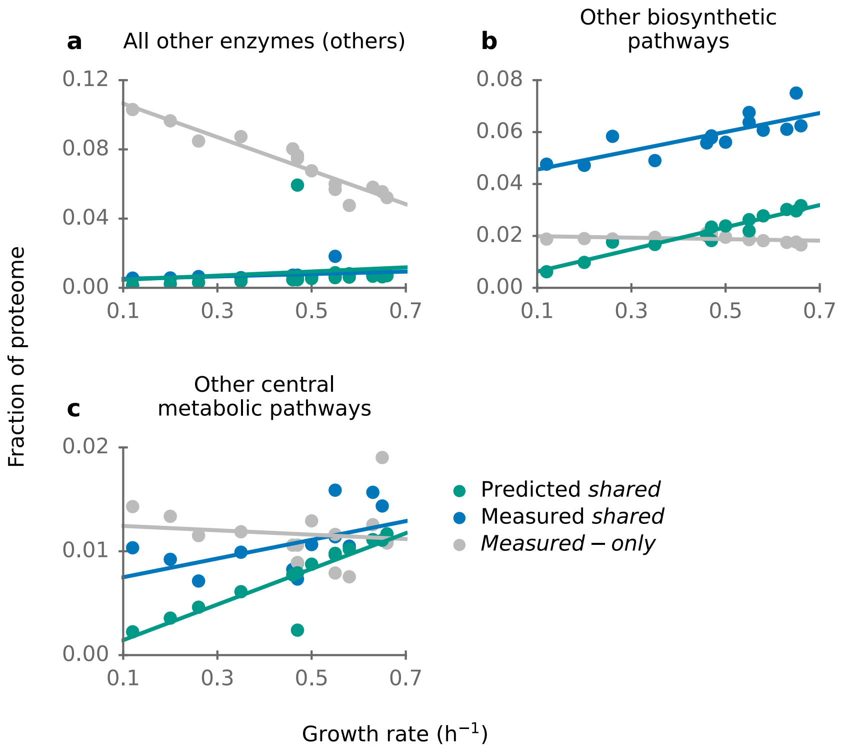


**Fig. S5. The proteome efficiency of other pathways.** (**a**) Combined proteome fraction of all enzymes in the *i*ML1515 model that are not part of the three pathway sets in Fig. 1b-1d ("Others"). While the abundance of those proteins predicted to be active under optimality is similar to the expected abundance ($GMFE_{pathway}$ = 1.79), a much higher proteome fraction is allocated to *measured-only* proteins. (**b**) Proteome investment into those biosynthetic proteins that are not part of the four pathway sets in Fig. 2. While predicted and observed concentrations of *shared* enzymes are strongly correlated ($r_{pathway}^2$ = 0.60; Table 1) and the expression of individual enzymes can be well explained by the predictions ($r_{individual}^2$ = 0.46), the corresponding values differ on average by almost 3-fold ($GMFE_{pathway}$ = 2.91). (**c**) Proteome investment into enzymes of central metabolism that are not part of the five pathway sets in Fig. 3. The prediction increases with growth rate, whereas the measured data is largely independent of growth rate.

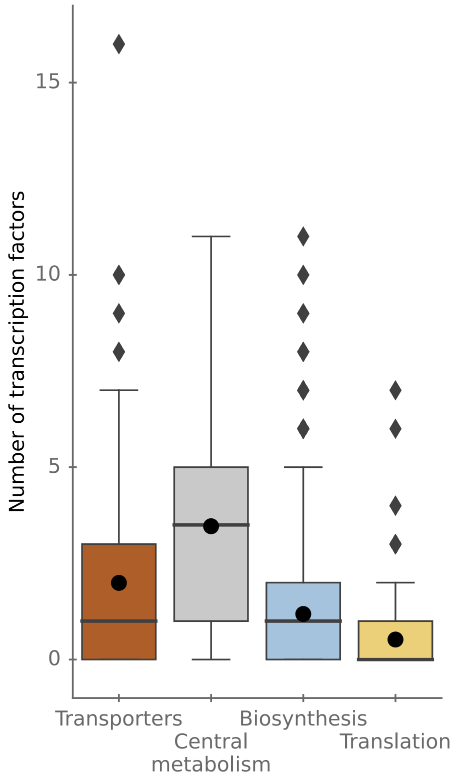


**Fig. S6. Genes in more central cellular processes are regulated by fewer transcription factors than genes in peripheral processes.** Genes encoding enzymes in biosynthesis pathways are regulated by fewer transcription factors than genes encoding transporters (one-sided Wilcoxon rank sum tests: $p < 10^{-10}$) and central metabolism ($p < 10^{-10}$). Genes in translation (here, all genes annotated with COG category "J"(4)) are also regulated by fewer transcription factors than genes in transporters ($p < 10^{-10}$) and central metabolism ($p < 10^{-10}$). Lines within boxes mark medians, dots indicate means. Boxes indicate 25% and 75% quantiles, whiskers extend from the boxes by 1.5x the interquartile range, diamonds are outliers.

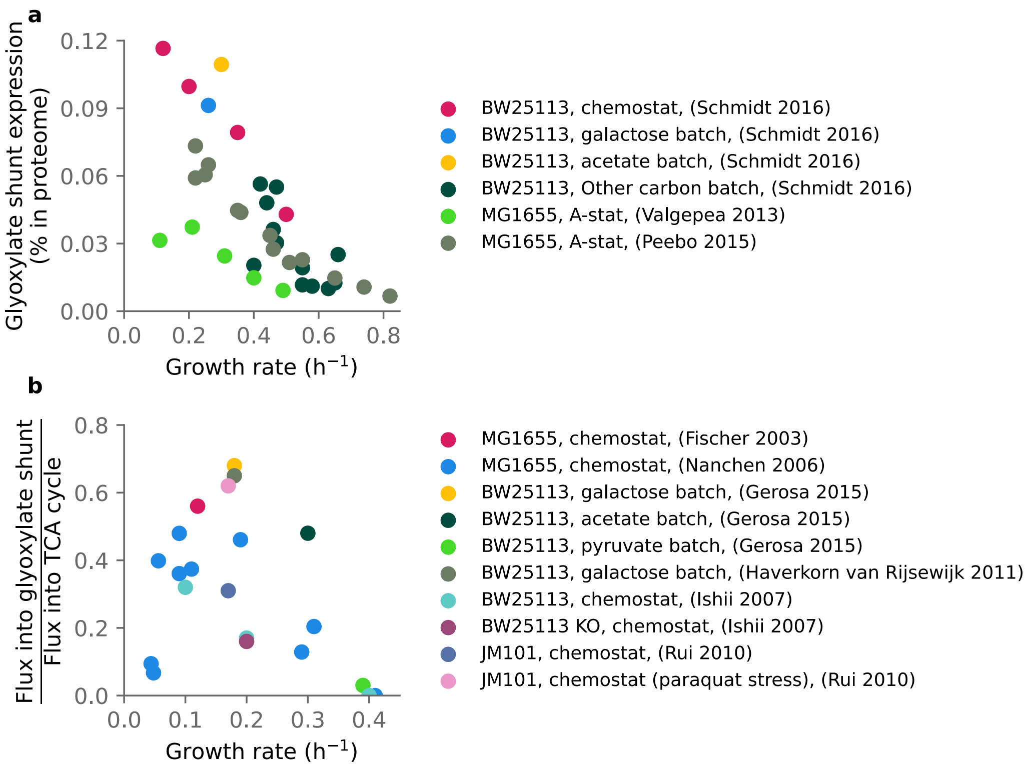


**Fig. S7. Growth rate-dependent activity of the glyoxylate shunt.** (**a**) Growth rate-dependent expression of the glyoxylate shunt, data from Refs. (5–7). (**b**) Growth rate-dependence of the flux through the glyoxylate shunt relative as a fraction of the flux into the TCA cycle. As the $^{13}$C metabolic flux analysis usually lumps linear sequential reaction sequences into a single reaction, flux into the TCA cycle is quantified through $^{13}$C measures of the flux of acetyl-CoA and oxaloacetate to isocitrate. The flux into the glyoxylate shunt is quantified through $^{13}$C measures of the flux catalyzed by Isocitrate lyase (isocitrate -> succinate + glyoxylate). Data from Refs. (8–13). Both protein abundance and flux decrease with increasing growth rate. The glyoxylate shunt's high protein abundance and large flux at low growth rates mark it as an important alternative to the TCA cycle in these conditions.
